# Uncovering directional epistasis in bi-parental populations using genomic data

**DOI:** 10.1101/2022.12.18.520958

**Authors:** Simon Rio, Alain Charcosset, Laurence Moreau, Tristan Mary-Huard

## Abstract

Epistasis, commonly defined as interaction effects between alleles of different loci, is an important genetic component of the variation of phenotypic traits in natural and breeding populations. In addition to its impact on variance, epistasis can also affect the expected performance of a population and is then referred to as directional epistasis. Before the advent of genomic data, the existence of epistasis (both directional and non-directional) was investigated based on complex and expensive mating schemes involving several generations evaluated for a trait of interest. In this study, we propose a methodology to detect the presence of epistasis based on simple inbred bi-parental populations, both genotyped and phenotyped, ideally along with their parents. Thanks to genomic data, parental proportions as well as shared parental proportions between inbred individuals can be estimated. They allow the evaluation of epistasis through a test of the expected performance for directional epistasis or the variance of genetic values. This methodology was applied to two large multi-parental populations, i.e., the American maize and soybean nested association mapping populations, evaluated for different traits. Results showed significant epistasis, especially for the test of directional epistasis, e.g., the increase in anthesis to silking interval observed in most maize inbred progenies or the decrease in grain yield observed in several soybean inbred progenies. In general, the effects detected suggested that shuffling allelic assocations of both elite parents had a detrimental effect on the performance of their progeny. This methodology is implemented in the EpiTest R-package and can be applied to any bi-/multi-parental inbred population evaluated for a trait of interest.

## Introduction

The term “epistasis” was first introduced by Bateson (1909) in the context of discrete traits with Mendelian segregation to describe the phenomena where the presence of an allele at one locus masks the effect of another locus. The extension of this concept to the context of populations of individuals evaluated for quantitative traits was presented by Fisher (1918) and is commonly referred to as “statistical epistasis”. In this context, epistasis characterizes statistical deviations between loci that arise after taking into account the effect of each locus independently (see Phillips (2008) for a review of the different views on epistasis).

Epistasis is prevalent in the genetic architecture of quantitative traits and it arises from the complex transcriptional, metabolic and biochemical networks involved in these traits (Kryazhimskiy, 2021). Despite its prevalence, epistasis has long been considered a nuisance parameter that can be ignored in breeding (Crow, 2010). This can be explained by the limited transmission of the epistatic part of the genotypic value between parents and offspring, as associations between alleles from different loci tend to be broken during meiosis. However, the contribution of epistasis to the genotypic value of a given commercial cultivar can be substantial, highlighting the importance of characterizing and predicting the epistatic component in plant breeding (Raffo et al., 2022; Varona et al., 2018).

To detect the presence of epistatic interactions, geneticists have designed complex crossing schemes, as summarized in Mather and Jinks (1982). These schemes required the production of several generations of progeny evaluated for the studied trait. The general principle consists of the comparison of the expected performance of the different generations allowing the isolation of epistatic terms whose significance can be tested. Examples include the well-known triple test cross (Jinks et al., 1969; Kearsey and Jinks, 1968) and other designs (Chahal and Jinks, 1978; Hayman, 1958; Melchinger, 1987). The ability of such approaches to detect epistasis relies on a set of non-cancelling epistatic genetic effects in the comparison of expected generation performances, leading to so-called directional epistasis (see Rouzic (2014) for a review).

An alternative for epistasis detection is to consider variance rather than expected performance. Cockerham (1954) and Kempthorne (1954) proposed a partitioning of the genetic variance into orthogonal components: additive, dominance, additive × additive, additive × dominance, dominance × dominance and higher order interactions. Since genetic variances are quadratic functions of genetic effects, the canceling of effects possibly observed in the expected performance is prevented. Before the use of molecular markers, the estimation of epistatic variances in heterozygous populations was very complex and reduced to a handful of sophisticated designs such as double cross hybrids (Rawlings and Cockerham, 1962a) or triallel analysis (Rawlings and Cockerham, 1962b). In the plant community, homozygous inbred individuals can be generated such as double-haploids (DH) or recombinant inbred lines (RILs). As the use of inbred progenies circumvents the difficulty of disentangling dominance from epistasis encountered in most designs, strategies were proposed to estimate additive and epistatic variance components. Choo (1981) proposed a total genetic variance partitioning into accross and within F2-derived DH populations, which can be solved to estimate additive and epistatic variance components. Other strategies were proposed based on diallel crosses (Choo, 1980) or random mating populations (Gallais, 1990).

The advent of molecular markers in the late 1980s revolutionized approaches to detect epistatic interactions. The identification of quantitative trait loci (QTL) involved in the genetic architecture of traits opened the way to the identification of epistasis using (i) one-dimensional approaches, i.e. by testing the interaction between a QTL and the genetic background (Blanc et al., 2006; Jannink and Jansen, 2001; Jannink et al., 2009), or (ii) two-dimensional approaches, i.e. by testing the interaction between pairs of QTL (Jannink et al., 2009; Kao et al., 1999). In the context of genome-wide associations studies, similar one-dimensional (Crawford et al., 2017; Jannink, 2007; Rio et al., 2020a) and two-dimensional (Hemani et al., 2011; Prabhu and Pe’er, 2012; Zhang et al., 2010) approaches were proposed. Molecular markers also make it possible to calculate genomic relationship matrices corresponding to the orthogonal partitioning of genetic variance (Álvarez-Castro and Carlborg, 2007; Vitezica et al., 2017), thus enabling to estimate epistatic variance components without the need for dedicated designs.

Inbred bi-parental populations, including Double Haploid (DH) or recombinant inbred lines (RIL) progenies, are pervasive in the plant genetics community. They are the cornerstone of breeding programs for self-pollinated species like wheat and soybean, as well as cross-pollinated species based on F1 hybrids between pure inbred lines like maize. They are also a fundamental component of genetic mapping studies based on single or multi-parental designs like nested association mapping (NAM) populations that have been generated for a large number of species over the last decade (Gage et al., 2020). With decreasing costs of genotyping, inbred bi-parental population datasets, including both genomic and phenotypic information, are becoming routinely available in most crops. In this study, we present a framework to test for the existence of epistasis in inbred bi-parental populations genotyped and evaluated for a trait, both through the expectation (i.e., directional epistasis) and the variance (i.e., non-directional epistasis) of genetic values. This framework is implemented in the new R-package “EpiTest” available from the CRAN. Applications are presented to two large NAM populations (the American maize NAM and the soybean NAM) evaluated for agronomy, phenology, morphology and/or quality traits.

## Material and Methods

### Infinitesimal model

To develop a procedure to test for the presence of epistasis, our approach consists in (i) introducing an infinitesimal model of genetic values of inbred bi-parental progeny with digenic epistatic interactions, (ii) deriving the expression of the expected genetic value and the covariance between genetic values, (iii) modeling phenotypic values through a Gaussian mixed model that inherits its design and covariance matrices from the expressions obtained in (ii), and (iv) proposing tests of model parameters involving only epistasis.

Let us consider a population of homozygous inbred individuals derived from a cross between two homozygous inbred parents A and B in absence of selection. Only polymorphic biallelic QTLs are considered with two genotypic states indicating both the genotype and the ancestry of alleles. Let *G*_*i*_ be the genetic value of individual *i*. One has:

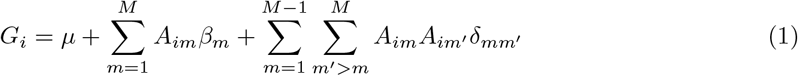

where *µ* is the genetic value of parent B, *M* is the number of loci, *A*_*im*_ is the allele ancestry indicator taking value 1 if individual *i* is homozygous for the parent A allele at locus *m* and 0 otherwise, *β*_*m*_ is the effect of substituting parent B allele by parent A allele at locus *m*, and *δ*_*mm*_*′* is the interaction effect generated by substituting parent B alleles by parent A alleles at loci *m* and *m*^*′*^. Note that this genetic model corresponds to the multi-linear model proposed by Hansen and Wagner (2001) to build the genotype-to-phenotype map for the population of interest, considering only digenic interactions between loci.

Let us assume a Bernoulli distribution for allele ancestries: *A*_*im*_ ∼ ℬ (*π*_*i*_), where *π*_*i*_ is the proportion of alleles originating from parent A for individual *i*, and cov (*A*_*im*_, *A*_*jm*_) = Δ_*ij*_ is the covariance between allele ancestries of individuals *i* and *j*. Note that allele ancestries are assumed to be independent between loci, which ignores the physical/genetic linkage between loci located on the same chromosome and constraints on allele ancestries due to a finite number of loci (see File S1 for details). The possible impact of the non-independence between allele ancestries on the method is addressed in the discussion. Both *π*_*i*_ and Δ_*ij*_ are key parameters to describe the genetic content of an individual (or a pair of individuals) in terms of (shared) parent ancestry proportions. While *π*_*i*_ may vary between 0 and 1, Δ_*ij*_ may vary between 0 and 0.25 when *i* = *j* and between 0.25 and -0.25 when *i* = *j*. These quantities can be used to compute expected genetic values and covariance between genetic values, which will later be used in a statistical model to test for epistasis, as described hereafter.

### Expectation and variance of genetic values

Let E (*G*_*i*_ |*π*_*i*_) be the expected genetic value conditional on the proportion of alleles originating from each parent. One shows that:

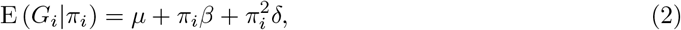

where 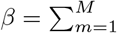 *β*_*m*_ is the linear “regression” coefficient of the genetic value on the parent A genome proportion and 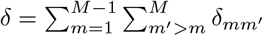 is the quadratic “regression” coefficient of the genetic value on the parent A genome proportion, which drives directional epistasis.

Similarly, let Cov(*G*_*i*_, *G*_*j*_ |Δ_*ij*_) be the covariance between genetic values conditional on shared proportion of alleles originating from each parent. One shows that:

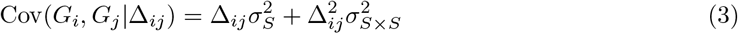

where 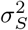 is the segregation variance generated by substituting parent B alleles by parent A alleles at loci over the whole genome and

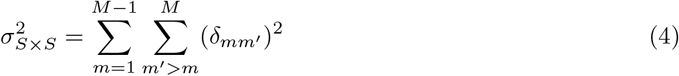

is the segregation × segregation (*S* ×*S*) interaction variance generated by the simultaneous substitution of parent B alleles by parent A alleles at loci pairs. Note that in a strict sense, 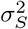 should be written 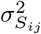 as it is a covariance parameter specific to the pair of individuals *i* and *j* that involves *π*_*i*_ and *π*_*j*_ in its expression. As the variation in 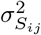 from a pair of individuals to another is expected to be marginal, except when the contribution of epistasis is high, it is in practice estimated as an overall parameter 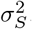. For the proofs of Eq. (2) and Eq. (3) as well as the complete expression of 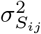, see File S1.

### Gaussian mixed model

The following model is assumed for phenotypic values:

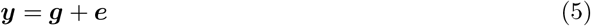

where ***y*** is the vector of reference phenotypes, e.g. best linear unbiased prediction (BLUP) or least-square mean calculated over the whole experimental design, ***g*** is the vector of genetic values defined below, ***e*** is the vector of errors with 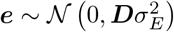, ***D*** is a diagonal matrix whose elements depend on the number of observations for each individuis 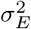 is the error variance, ***g*** and ***e*** are independent.

Regarding ***g***, one can derive from the infinitesimal model in Eq. (1) an approximate Gaussian variance component model that inherits its mean and variance components from Eq. (2) and Eq. (3). The genetic values are then modeled as the sum of a fixed intercept and two random components independent from each other:

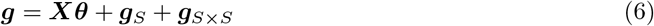

where ***X*** = (**1, *π, π***^2^) is the design matrix for fixed effects with ***π*** being the vector of parent A proportions, ***θ*** = (*µ, β, δ*)^*T*^ being the vector of fixed effects, ***g***_*S*_ is the vector of the segregation component of the genetic value, and ***g***_*S S*_ is the vector of the *S* _*×*_ *S* interaction component of the genetic value. Each random genetic effect is modeled as being drawn from a normal distribution with a covariance structure inherited from the infinitesimal model: 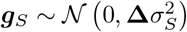 and 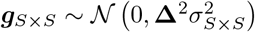, where **Δ** is the matrix of coefficients Δ_*ij*_.

### Inference and tests

In practice, ***π*** can be estimated using:

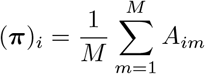

and **Δ** using:

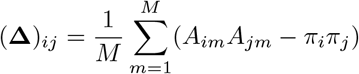

Based on Eq. (5) and Eq. (6), two tests can be proposed whose respective null hypotheses are:

- *H*_0_ : *δ* = 0 to test the existence of directional epistasis in the fixed part of the model,
- *H*_0_: 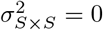 to test the contribution of epistasis to the genetic variance.

As **Δ** and **Δ**^2^ diagonal elements may vary between 0 and 0.25 or 0 and 0.0625, respectively, their corresponding variance component are not scaled to the phenotypic variance. To ease the comparison of variances, all covariance matrices (**Δ, Δ**^2^ and ***D***) are standardized as in Vitezica et al. (2017) by dividing each matrix by a scalar so that the mean of diagonal elements equals 1. The scalar corresponds to the sum of diagonal elements divided by the dimension of the square matrix. The inference of fixed and variance parameters is done using restricted maximum likelihood (ReML). The nullity of the *δ* parameter (i.e., directional epistasis test) can be tested using a Wald test for which the statistics has a chi-squared distribution. The test involving the variance parameter 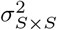 is based on a likelihood ratio test for which the distribution of the statistics is a specific mixture of chi-square distributions (Self and Liang, 1987; Stram and Lee, 1994). In what follows, all tests were adjusted for multiple testing using a 5% Bonferroni threshold over the number of populations in each dataset.

### Datasets

Two NAM populations were considered in this study, each including multiple bi-parental populations evaluated for several traits of interest.

The soybean NAM is described in Song et al. (2017). It consists of 39 families of approximately 140 recombinant inbred lines (see Table S1 for exact numbers) generated by crossing 39 diverse inbred lines to the central inbred line “IA3023”. All individuals were genotyped using the “SoyNAM6K BeadChip” for which the reference allele (coded 0) corresponds to the homozygous genotype for “IA3023” alleles and the alternative allele (coded 1) corresponds to the homozygous alleles for the other parent. The number of polymorphic markers ranged from 2,464 to 3,783 according to the population (mean and standard deviation of 3,245 and 432, respectively). As described in Diers et al. (2018) and Xavier et al. (2018), the NAM populations were evaluated along with parental lines in 19 trials for grain yield in kg/ha, days to maturity, plant height in centimeter, lodging score from 1 to 5, grain moisture in percentage of humidity, and protein/oil/fiber in percentage of the grain content. The BLUPs of genotypic performances were calculated over the whole design using a Gaussian model including a random genotype effect and a random trial effect fitted using the “lme4” R-package (Bates et al., 2015).

The American maize NAM is described in Yu et al. (2008) and McMullen et al. (2009). It consists of 25 families of approximately 200 recombinant inbred lines (see Table S2 for exact numbers) generated by crossing 25 temperate and tropical inbred lines to the central inbred line “B73”. All individuals were genotyped for 1,106 SNPs polymorphic in all populations (Buckler et al., 2009) for which the reference allele (coded 0) corresponds to the homozygous genotype for “B73” alleles and the alternative allele (coded 1) corresponds to the homozygous alleles shared by all parental lines but B73 (i.e., only SNPs involving alleles specific to B73 were considered). The NAM populations were evaluated along with parental lines in ten trials (combinations of five locations and four years) for days to anthesis/silking and anthesis to silking interval in growing degree days, plant height and ear height in centimeter. The BLUPs of genotypic performances over the whole design used in this study were those calculated by Peiffer et al. (2014).

The model in Eq. (6) was applied separately to all maize and soybean NAM populations, where parent A refers to the central parent and parent B refers to the alternative parent. For all datasets, a single reference phenotype was considered for each individual after adjusting for experimental design effects. In both the maize and the soybean experiments, parental lines were used as checks and thus had a much larger number of observations, leading to a better precision associated with the resulting reference phenotype. The diagonal elements of ***D*** from Eq. (6) were calculated accordingly and are summarized in Table 1.

**Table 1:**
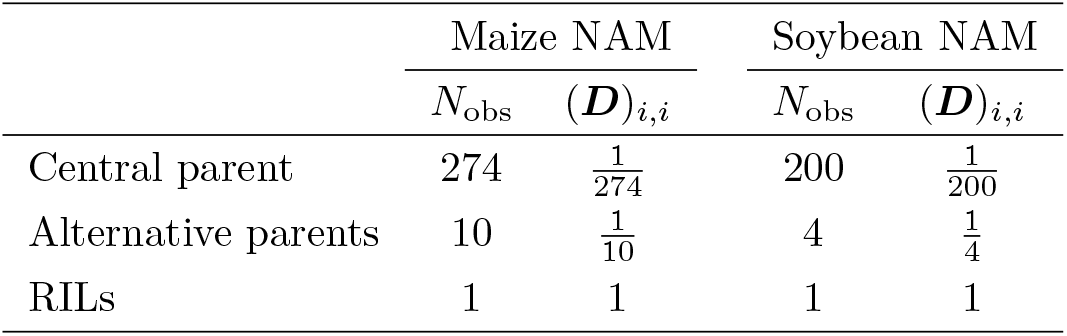
Number of observations for parental lines used as checks relative to that of RILs (*N*_obs_) and corresponding diagonal elements of the error covariance matrix (***D***)_*i,i*_

|  | Maize NAM |  | Soybean NAM |  |
| --- | --- | --- | --- | --- |
| | $N_{\text{obs}}$ | $(\mathbf{D})_{i,i}$ | $N_{\text{obs}}$ | $(\mathbf{D})_{i,i}$ |
| Central parent | 274 | $\frac{1}{274}$ | 200 | $\frac{1}{200}$ |
| Alternative parents | 10 | $\frac{1}{10}$ | 4 | $\frac{1}{4}$ |
| RILs | 1 | 1 | 1 | 1 |

### Simulation procedure summary

The precision of parameter estimates as well as the false positive rate and the statistical power of epistasic tests were evaluated with simulated traits using the American maize NAM genotypic data (see File S2 for details). Two genetic scenarios were defined and simulated 100 times: “No-epistasis” where only additive effects were simulated and “Epistasis” where both additive and epistatic effects were simulated. For the “Epistasis” genetic scenario, the simulated epistatic variance was relatively homogeneous across simulations while the simulated directional epistasis was either positive or negative and could take a value close to 0 up to a large deviation from 0. This is because the directionality of epistasis was not directly controlled in the simulations but was generated by the randomness of simulated QTL effects and evaluated afterwards. From these simulated genetic architectures, traits were simulated with different heritabilities *h*^2^ ∈ {0.25, 0.5, 0.75} based on a design with different population sizes *N* ∈ {60, 90, 120, 150, 180}.

### Data availability statement

The test procedure is implemented in a new R-package “EpiTest”, which is based on the mixed model inference R-package “MM4LMM” (Laporte et al., 2022), both available from the CRAN. The American maize NAM dataset is available at: https://www.panzea.org/. The phenotypic file considered was: “Peiffer2014Genetics_blupPhenos20150325.xlsx” and the genotypic file was: “NAM_genos_imputed_20090807.xlsx”. The soybean NAM dataset is available at https://www.soybase.org/SoyNAM/. The phenotypic file considered was the file: “SoyNAM all familes phenotype data.txt” and the genotypic file considered was the file: “SoyNAM parents+progeny 4312 SNP genotypes Wm82.a2”.

## Results

### Simulation results summary

Using simulations (see File S2 for details), the ability of the model to estimate parameters accurately was confirmed. Both tests effectively controlled for false positives at the required nominal level of 5% for different population sizes and heritabilities, with a maximum observed proportion of false positives of 6.67% for the directional epistasis test when *N* = 60 and *h*^2^ = 0.75 and a maximum of 5.79% for the epistatic variance test when *N* = 120 and *h*^2^ = 0.50 (File S2). For the variance test, the observed proportion of false positives could be substantially lower than expected when the population size dropped to 60 individuals (minimum of 1.17% instead of 5% when *h*^2^ = 0.75). As expected, for both tests, the statistical power increased with increasing population size and heritability. The statistical power of the epistatic variance test was generally very low, except for large populations with high *h*^2^ where the power was moderate (Figure 1**B**). For the directional epistasis test, the statistical power could be very high (i.e. close 100%) for the simulated genetic architectures that generated a large |*δ*|. In summary, the directional epistasis test appears more powerful than the epistatic variance test, provided that *δ* substantially deviates from 0.

**Figure 1:**
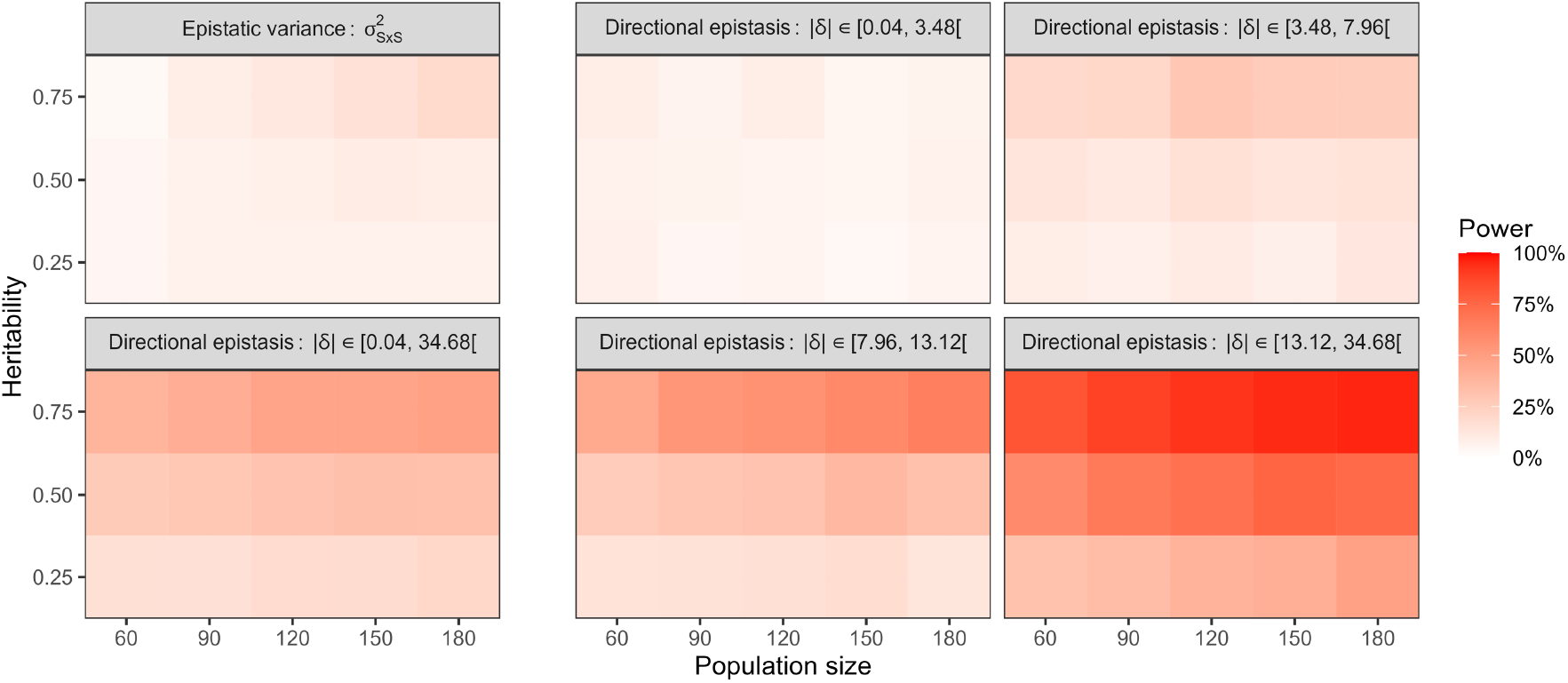
Statistical power of the epistatic variance and directional epistasis tests evaluated based on the American maize NAM populations and 100 genetic architectures in presence of epistasis. Statistical power was calculated as the proportion of significant tests (*α* = 5%) and was evaluated for different population sizes (*N*) and heritabilities (*h*^2^). For the directional epistasis parameter *δ*, simulated genetic architectures were considered either globally (|*δ*| ∈ [0.04, 34.68[) or clustered into quartiles according to the absolute value of the simulated *δ*: the first quartile with |*δ*| ∈ [0.04, 3.48[, the second quartile with |*δ*| ∈ [3.48, 7.96[, the third quartile with |*δ*| ∈ [7.96, 13.12[and the last quartile with |*δ*| ∈ [13.12, 34.68[.

The superiority in statistical power of the directional epistasis test over the epistatic variance test was also confirmed by the total number of significant tests found for real data (Table 2), with 99 and 12 significant tests for the directional epistasis and epistatic variance tests, respectively.

**Table 2:**
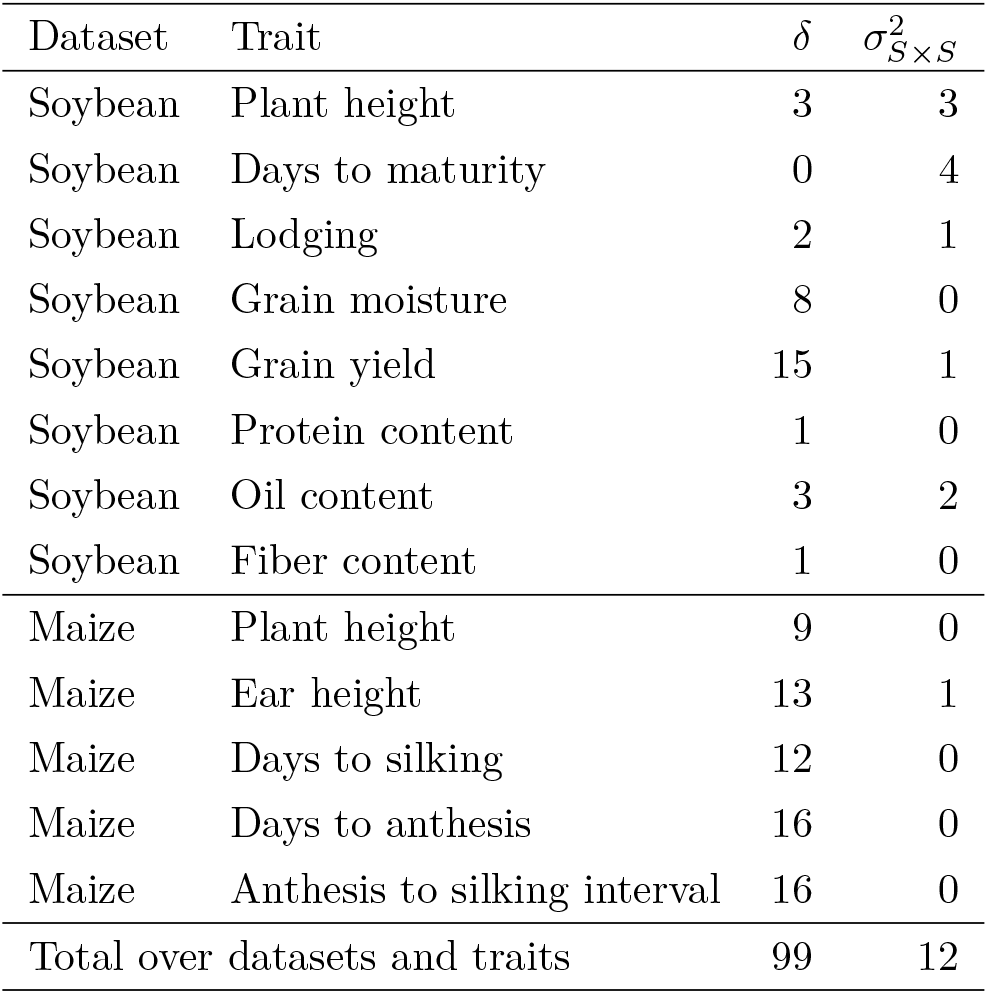
Number of significant tests based on a Bonferroni threshold at a nominal level of 5% over the number of populations for the directional epistasis test (*δ*) and the epistatic variance test 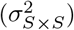 using the model of Eq. (6)

| Dataset | Trait | $\delta$ | $\sigma_{S \times S}^2$ |
| --- | --- | --- | --- |
| Soybean | Plant height | 3 | 3 |
| Soybean | Days to maturity | 0 | 4 |
| Soybean | Lodging | 2 | 1 |
| Soybean | Grain moisture | 8 | 0 |
| Soybean | Grain yield | 15 | 1 |
| Soybean | Protein content | 1 | 0 |
| Soybean | Oil content | 3 | 2 |
| Soybean | Fiber content | 1 | 0 |
| Maize | Plant height | 9 | 0 |
| Maize | Ear height | 13 | 1 |
| Maize | Days to silking | 12 | 0 |
| Maize | Days to anthesis | 16 | 0 |
| Maize | Anthesis to silking interval | 16 | 0 |
| Total over datasets and traits |  | 99 | 12 |

### Soybean NAM

The method was first applied to the soybean NAM that includes 39 bi-parental families evaluated for grain yield, days to maturity, plant height, lodging, grain moisture, and protein/oil/fiber content.

The presence of directional epistasis was investigated using the directional epistasis test (Table 2, see the Table S1 for details on model parameters estimates and tests). A focus was done on plant height (Figure 2B), grain yield (Figure 2D), and oil content (Figure 2F), results for the other traits are shown in supplementary material (Figures S1B, S2B, S3B, S4B and S5B). Significant directional epistasis was found for 3, 15 and 3 populations out of 39 for plant height, grain yield and oil content, respectively. For grain yield, significant quadratic coefficients *δ* were all positive, implying that progenies had lower average performances than expected under a model without directional epistasis. Using model parameter estimates presented in the Table S1, expected performances can be compared between parents and progeny by considering a reference progeny with perfectly balanced parental ancestry proportions of *π*_*i*_ = 0.5. For instance, for the population IA3023 × TN05.3027, IA3023 has an expected performance of 3937.2 kg/ha, TN05.3027 has an expected performance of 3892.3 kg/ha, and the expected performance of their progeny (with perfectly balanced ancestry proportions) is below that of the two parents: 3640.7 kg/ha, indicating the presence of directional epistasis.

**Figure 2:**
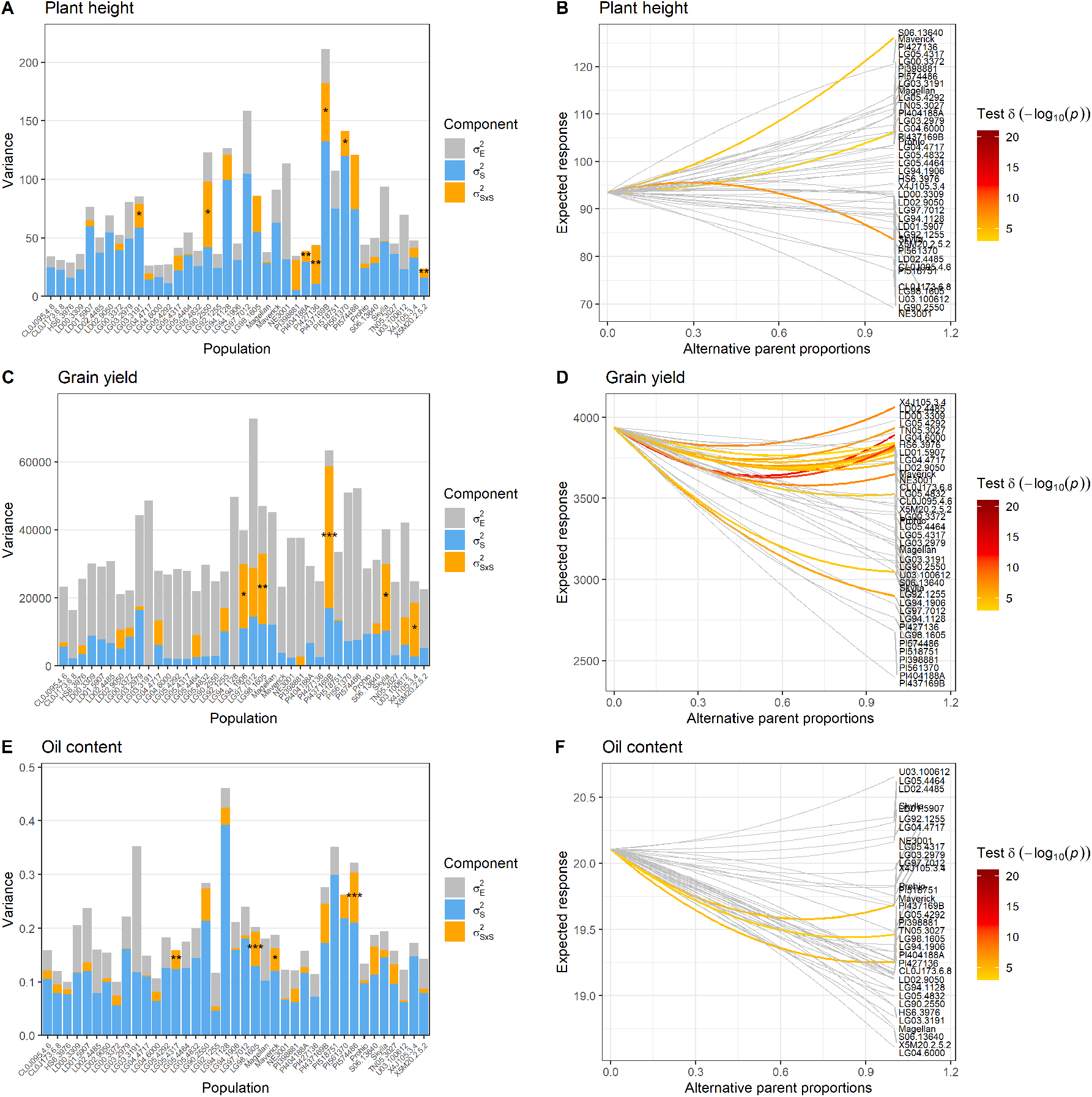
Variance component barplots and directional epistasis plots of the soybean NAM for three traits: plant height (**A** and **B**), grain yield (**C** and **D**) and oil content (**E** and **F**). For the variance components barplots, the range of p-value obtained for the likelihood ratio test of 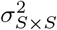 are indicated as: * for p-values inferior to 0.05, ** for p-values inferior to 0.01, and *** for p-values inferior to 0.001. For the directional epistasis plots, tests with a p-value higher than the Bonferroni threshold at a nominal level of 5% over the number of populations are indicated in grey.

The contribution of epistasis to the genetic variance in the progeny was investigated using the test of the variance part of the model (Table 2, see Table S1 for details on model parameters estimates and tests). A focus was done on the same traits as for directional epistasis (see Figures 2A, 2C and 2E), results for the other traits are shown in supplementary material (Figures S1A, S2A, S3A, S4A and S5A). In contrast to the directional epistasis tests, only a few epistatic variance tests were significant. Most traits showed a limited contribution of the epistatic variance component 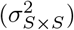 to the genetic variance, which might be caused by the limited precision in estimating this parameter. Notable exceptions include the IA3023×PI437169B population for grain yield.

### American maize NAM

The method was then applied to the American maize NAM that includes 25 bi-parental families evaluated for days to anthesis/silking, anthesis to silking interval, and ear/plant height.

The presence of directional epistasis was investigated using the directional epistasis test (Table 2, see Table S1 for details on model parameters estimates and tests). A focus was done on days to anthesis (Figure 3B), anthesis to silking interval (Figure 3D), and plant height (Figure 3F), results for days to silking and ear height are shown in supplementary material (Figure S6B and Figure S7B, respectively). Significant directional epistasis was found for 16, 16 and 9 populations out of 25 for days to anthesis, anthesis to silking interval and plant height, respectively. For anthesis to silking interval, all significant *δ* coefficients were negative, suggesting that progenies had higher average values than expected under a model without directional epistasis.

**Figure 3:**
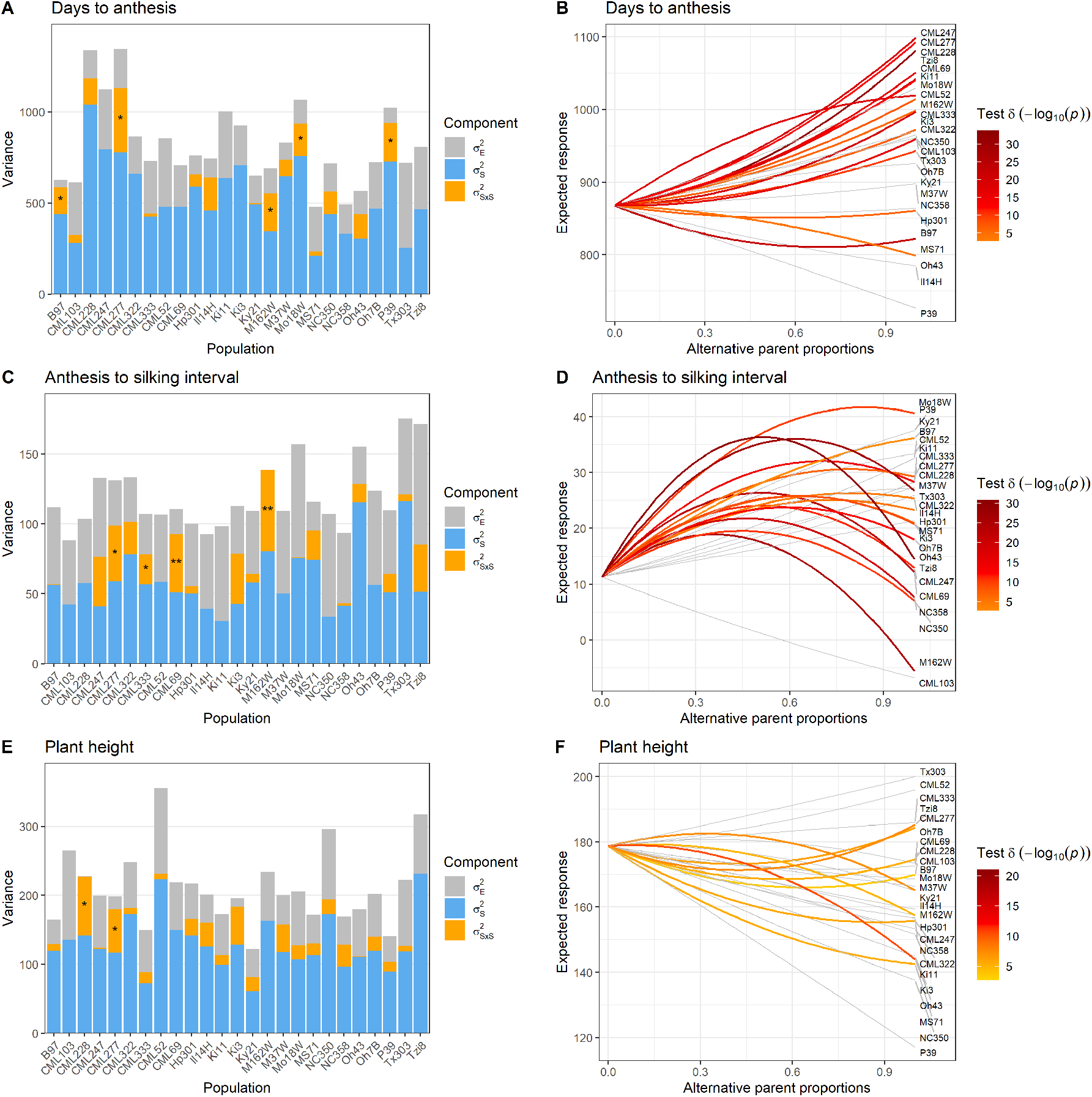
Variance component barplots and directional epistasis plots of the American maize NAM for three traits: days to anthesis (**A** and **B**), anthesis to silking interval (**C** and **D**) and plant height (**E** and **F**). For the variance components barplots, the range of p-value obtained for the likelihood ratio test of 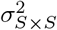 are indicated as: * for p-values inferior to 0.05, ** for p-values inferior to 0.01, and *** for p-values inferior to 0.001. For the directional epistasis plots, tests with a p-value higher than the Bonferroni threshold at a nominal level of 5% over the number of populations are indicated in grey

The contribution of epistasis to the genetic variance in the progeny was investigated using the test of the variance part of the model (Table 2, see Table S2 for details on model parameters estimates and tests). A focus was done on the same traits as for directional epistasis (see Figures 3A, 3C and 3E). The same figures for other traits are presented in Figures S6A and S7A for days to silking and ear height, respectively. In contrast to the directional epistasis tests, only a few epistatic variance tests were significant. Most traits showed a limited to moderate contribution of the epistatic variance component 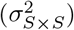 to the genetic variance.

## Discussion

### Epistasis tests for inbred bi-parental populations

We developed a procedure to test for the presence of epistasis in inbred bi-parental progeny when genomic information is available. The procedure is based on a Gaussian mixed model where epistasis is accounted for through a mean component (*δ*) and a variance component 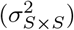.

A significant *δ* for the trait indicates non-cancelling epistatic effects and can be defined as directional epistasis (Rouzic, 2014). In genomic prediction models for outbred populations, marker effects are usually assumed to be centered around zero. While this assumption holds true for additive effects, one expects to observe mostly positive dominance effects especially for traits showing heterosis and thus a directional dominance. Xiang et al. (2016) proposed the use of an inbreeding covariate to account for directional dominance in genomic prediction models. The concept of directional epistasis presented in our study is similar to the latter in that it involves non-cancelling interaction effects over loci pairs. It would be interesting to see to what extent the simultaneous consideration of parental proportions and inbreeding in the modeling of non-homozygous bi-parental populations (e.g. F2) would help disentangling the effect of directional dominance and epistasis. Directional epistasis is expected not only in bi-parental populations, but also in any type of population. We plan to investigate how to extend our model to account for the directionality of epistasis in genomic prediction models in the near future.

The two parameters *δ* and 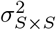 only involve epistatic effects and consequently will have value 0 whenever these effects are null. However, *δ* can be zero in the presence of epistatic interactions, provided that the effects cancel each other out over loci pairs. Also note that the structure of the covariance matrix involves a covariance Δ_*ij*_ between parental allele ancestries that has already been presented in the context of admixed populations (Aase et al., 2022; Rio et al., 2020b) for the segregation of group-specific allele ancestries. The use of a squared genomic relationship matrix (i.e, 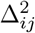 in our study) to estimate the additive×additive epistatic variance component has also been suggested in the context of outbred populations (Vitezica et al., 2017).

In absence of epistasis, the expected genetic value of an inbred offspring with perfectly balanced parental ancestry proportions does not deviate from the average value of its inbred parents. The genetic variance is then only driven by the effect of segregating QTL parental alleles. When epistasis is present, the expected genetic value can deviate from the average value of its inbred parents, and is also accompanied with an additional source of variability generated by the simultaneous segregation of QTL parental alleles at loci pairs.

The proposed test procedure requires the genomic information to be available in order to calculate the covariance matrix between allele ancestries associated with the variances 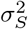 and 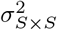. In absence of genomic information, a test could theoretically be implemented using a linear model that considers expected parental proportions of 0.5 for the whole bi-parental population, provided that both parents are also evaluated. Note that it is not possible to distinguish between the two genetic variances and the model is no longer identifiable whenever the genetic information is not available. As a consequence, the epistatic variance 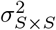 cannot be tested. Based on simulation, the directional epistasis test prove to be more powerful than the test of the epistatic variance (Figure 1 and File S2). This observation is consistent with the larger number of significant directional epistasis tests compared to the test of epistatic variance for real traits in both NAM datasets (Table 2). However, some traits such as days to maturity in soybean have a higher number of significant tests for 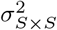 than for *δ*, which demonstrates the benefit of testing both parameters to detect epistasis. Note that, thanks to genomic information, the test procedure can be applied only based on the progeny data, i.e. in absence of any parental phenotypic information. While this may be useful in some specific cases, we recommend experimental designs that include a thorough evaluation of both parents since the parental information is expected to have a strong (positive) impact on the estimation of model parameters. In practice, parents are often evaluated along with their progeny as replicated checks in bi- and multi-parental experimental designs - as in the maize and soybean NAM datasets - which provides favorable conditions for the application of the proposed procedure.

To make the derivation of the model feasible, we made the common assumption of independence between QTL. However, this assumption is violated in segregating bi-parental populations due to genetic linkage between loci on the same chromosomes. Regarding the properties of statistical tests, we showed that they respected the nominal level of 5% type I error rate when applied to simulated traits in presence of genetic linkage between markers. The impact of genetic linkage on the value of genetic parameters is more difficult to evaluate. In presence of non-independent QTLs, the expression of the epistatic variance would be different from the simple sum of squares of interaction effects in Eq. (4) obtained when assuming independence. In the case of two perfectly co-segregating QTLs in the population, the two QTL effects and their interaction effect would be mixed into a single segregation effect of the QTL block. In our simulations, we estimated the fixed and variance parameters in presence of linked markers and compared them to the ones calculated using the true simulated effects. No strong bias was observed when comparing the estimates to the calculated values. (File S2).

In this study, only epistatic interactions between pairs of loci were considered. We believe that this constraint is reasonable as the contribution of epistatic interaction to the fitness landscape is expected to decrease with increasing order of interaction (Weinreich et al., 2018). However, this approach could theoretically be generalized to higher order interactions by increasing the power to which the parental proportions in the fixed part of the model and the covariance matrix between allele ancestries are raised. One should keep in mind that this would be done at the cost of increased model complexity and the statistical power to detect such higher order effects would likely be low. Regarding variance components, the comparison of the range of coefficients of **Δ** and **Δ**^2^ (e.g. in the bi-parental population B73 × B97 in Fig. S8) shows that the off-diagonal elements of **Δ**^2^ are shrunk toward 0. We can expect this phenomenon to be even more pronounced for higher order interactions.

The strength of the test procedure presented in this study lies in its ability to target epistasis both through the expectation and the variance of genetic values without requiring a complex design, as is usually the case (Mather and Jinks, 1982). Only simple inbred progenies (e.g., DH or RIL progeny) need to be genotyped and evaluated for a trait of interest. Such datasets are already available in large numbers and can simply be recycled to investigate the presence of epistasis. Our procedure also opens the way to the systematic investigation of epistasis in plant breeding programs and quantitative genetics studies. We highly recommend applying this test procedure prior to any QTL analysis as it helps determine the need to test for pairwise epistatic QTL effects.

### Epistasis in breeding

Epistasis has long been considered a nuisance parameter that can be ignored in breeding (Crow, 2010). However, its presence can influence both the average performance of a cross, through the phenomenon of directional epistasis (Rouzic, 2014), and the genetic variance generated by that cross, through an additional epistatic variance component. The average performance and the genetic variance of a cross are the two parameters involved in the usefulness criterion proposed by Schnell and Utz (1975). A good understanding of the genetic determinism of traits subject to selection should allow a better reasoning of breeding mating designs, i.e. choosing which parents to cross along with the minimum progeny size that needs to be generated to maximize the chances of obtaining a superior individual for the trait.

In maize, several experiments have been conducted in which directional epistasis was detected for grain yield, forage yield, plant height, ear height, kernel row number, maturity or flowering traits (Bauman, 1959; Gamble, 1962; Hallauer and Russell, 1962; Melchinger et al., 1986, 1988; Wolf and Hallauer, 1997), or through epistatic variance for grain yield, forage yield and grain dry matter content (Melchinger et al., 1988). Similarly, significant epistasis has also been detected in soybean through variance for oil content (Hanson and Weber, 1961) or through directional epistasis for grain yield (Barona et al., 2012; Uzokwe et al., 2017). The results of these experiments are consistent with the results of our study demonstrating the prevalence of epistasis affecting the mean and variance of breeding traits in maize and soybean.

For some traits like anthesis to silking interval in maize or grain yield in soybean, the significant directional epistasis detected is always accompanied with a deterioration of the genetic values compared to that expected based on parental values under a purely additive model. Regarding maize, anthesis to silking interval increases in most progenies. This desynchronization of male and female flowering is a well-known indicator of stress (Edmeades et al., 2015) and is not a desirable trait in maize breeding. In the case of soybean, a decrease in grain yield is non-desirable as it directly conditions farmers’ income. In general, it is reasonable to assume that a good elite inbred line results from both a good additive genetic value and a combination of alleles with favorable epistatic effects. When crossing two unrelated elite inbred lines, favorable allele combinations are likely to be broken, thus leading to deterioration of the progeny mean. A similar perspective is that of less-than-additive epistasis (Eshed and Zamir, 1996), where the effect of QTL becomes low in a genetic background with many favorable alleles. These hypotheses are supported by the tendency for directional epistasis to occur between parents with high grain yield in soybean.

A major challenge remains to understand what other factors determine the apparition of directional epistasis for a given cross. This question is particularly relevant for traits like plant height in maize displaying directional epistasis with opposite sign depending on the bi-parental population. One may hypothesize to observe more directionnal epistasis when crossing distant lines but our results do not completely support it. For instance, tropical maize inbred lines (e.g., lines with names starting with CML) do not systematically generate the largest directional epistasis when crossed to the distant stiff-stalk parent B73. A better solution to predict the apparition of directional epistasis would probably be a fine characterization of the genetic architecture of traits, including the number and position of QTLs along with their marginal and combined effects.

## Notes

https://www.panzea.org/

https://www.soybase.org/SoyNAM/

### Competing Interest Statement

The authors have declared no competing interest.

### Summary of Updates

Changes in figures and text

