## Supplementary material for "Uncovering directional epistasis in bi-parental populations using genomic data": File S1

### File S1: Statistical theory

Let us consider a population of homozygous inbred individuals derived from a cross between two homozygous inbred parents A and B. Only polymorphic biallelic QTLs are considered with two genotypic states indicating both the genotype and the ancestry of alleles: homozygous for parent A or parent B alleles. In this context, the genetic value of individuals for a trait of interest can be modeled as:

$$G_i = \sum_{k \in S} \sum_{m=1}^{M} A_{imk} \beta_{mk} + \sum_{k \in S} \sum_{k' \in S} \sum_{m=1}^{M-1} \sum_{m'>m}^{M} A_{imk} A_{im'k'} \delta_{mm'kk'} \tag{1}$$

where:

- $G_i$: genetic value of individual $i$
- $S = \{A, B\}$: set of parents
- $M$: number of QTL
- $A_{imk}$: allele ancestry indicator taking the value 1 if individual $i$ is homozygous for parent $k$ alleles at locus $m$, and 0 otherwise
- $A_{imB} = 1 - A_{imA}$
- $\beta_{mk}$: main QTL allele effects that are associated with a homozygous genotype for parent $k$ alleles at locus $m$
- $\delta_{mm'kk'}$: deviation QTL allele effects that are associated with the homozygous genotype for parent $k$ alleles at locus $m$ and for parent $k'$ at locus $m'$

Note that the model in Eq. (1) can be equivalently written using one of the parents as a reference (here parent B) using:

$$G_i = \mu + \sum_{m=1}^{M} A_{imA} \beta_m + \sum_{m=1}^{M-1} \sum_{m'>m}^{M} A_{imA} A_{im'A} \delta_{mm'} \tag{2}$$

where

$$\mu = \sum_{m=1}^{M} \beta_{mB} + \sum_{m=1}^{M-1} \sum_{m'>m}^{M} \delta_{mm'BB} \tag{3}$$

is the genetic value of parent B,

$$\beta_m = \left( \beta_{mA} + \sum_{m \neq m'}^{M} \delta_{mm'AB} \right) - \left( \beta_{mB} + \sum_{m \neq m'}^{M} \delta_{mm'BB} \right) \tag{4}$$

is the effect of substituting parent B alleles by parent A alleles at locus $m$ in a parent B genetic background, and

$$\delta_{mm'} = (\delta_{mm'AA} - \delta_{mm'BA}) - (\delta_{mm'AB} - \delta_{mm'BB}) \tag{5}$$

is the interaction effect generated by substituting parent B alleles by parent A alleles at locus $m$ and $m'$.

#### Model assumptions

The following assumptions are considered for the model in Eq. (1):

- $A_{imk} \sim \mathcal{B}(\pi_{ik})$: a Bernoulli distribution is assumed for allele ancestries $A_{imk}$ with parameter $\pi_{ik}$
- $\pi_{ik}$: proportion of alleles originating from parent $k$ for individual $i$
- $\pi_{iB} = 1 - \pi_{iA}$
- $\mathrm{E}\left(A_{imk} A_{jmk}\right) = \theta_{ij}^{k}$: shared ancestry proportions for parent $k$ alleles between individuals $i$ and $j$ (see Fig. A)
- $\theta_{ij}^{B} = 1 - \pi_{iA} - \pi_{jB} + \theta_{ij}^{A}$
- $\mathrm{Cov}\left(A_{imk}, A_{jmk}\right) = \theta_{ij}^{k} - \pi_{ik}\pi_{jk} = \Delta_{ij}$: covariance between allele ancestries
- $\mathrm{Cov}\left(A_{imk}, A_{jm'k'}\right) = 0 \; \forall i, j, m \neq m', k, k'$: allele ancestries are assumed to be independent between loci, which is a simplifying assumption ignoring the physical/genetic linkage between loci located on the same chromosome and constraints on allele ancestries due to a finite number of loci (e.g., when $M = 2$ and $\mathrm{E}\left(A_{imA}\right) = \pi_{iA} = 0.5$, $\mathrm{E}\left(A_{imA} | A_{im'A} = 1\right) = 0 \neq \mathrm{E}\left(A_{imA}\right)$)

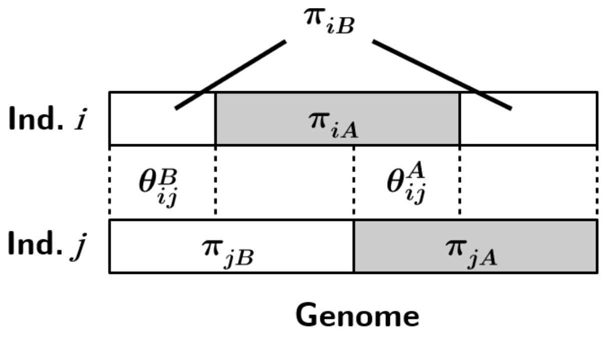


Figure A: Diagram illustrating the genome-wide allele ancestry of two inbred individuals $i$ and $j$, represented by their haplotypes, with a proportion $\pi_{iA}$ ($\pi_{iB}$) of genome A (B) for $i$ and $\pi_{jA}$ ($\pi_{jB}$) of genome A (B) for $j$, and a proportion of shared group A (B) ancestry $\theta_{ij}^{A}$ ($\theta_{ij}^{B}$) between $i$ and $j$. Note the existence of the following constraint: $\theta_{ij}^{B} = 1 - \pi_{iA} - \pi_{jA} + \theta_{ij}^{A}$

#### Expected genetic value

Let $\mathrm{E}\left(G_i|\pi_{ik}\right)$ be the expected genetic value conditional on the proportion of alleles originating from each parent:

$$\mathrm{E}\left(G_i|\pi_{ik}\right) = \mathrm{E}\left(\sum_{k\in S}\sum_{m=1}^{M} A_{imk}\beta_{mk} + \sum_{k\in S}\sum_{k'\in S}\sum_{m=1}^{M-1}\sum_{m'>m}^{M} A_{imk}A_{im'k'}\delta_{mm'kk'}\right) \tag{6}$$

$$\mathrm{E}\left(G_i|\pi_{ik}\right) = \sum_{k\in S}\sum_{m=1}^{M} \mathrm{E}\left(A_{imk}\right)\beta_{mk} + \sum_{k\in S}\sum_{k\in S}\sum_{m=1}^{M-1}\sum_{m'>m}^{M} \mathrm{E}\left(A_{imk}A_{im'k'}\right)\delta_{mm'kk'} \tag{7}$$

$$\mathrm{E}\left(G_i|\pi_{ik}\right) = \sum_{k\in S}\pi_{ik}\sum_{m=1}^{M}\beta_{mk} + \sum_{k\in S}\sum_{k'\in S}\pi_{ik}\pi_{ik'}\sum_{m=1}^{M-1}\sum_{m'>m}^{M}\delta_{mm'kk'} \tag{8}$$

$$\mathrm{E}\left(G_i|\pi_{ik}\right) = \mu + \pi_{iA}\sum_{m=1}^{M}\beta_m + \pi_{iA}^2\sum_{m=1}^{M-1}\sum_{m'>m}^{M}\delta_{mm'} \tag{9}$$

$$\mathrm{E}\left(G_i|\pi_{ik}\right) = \mu + \pi_{iA}\beta + \pi_{iA}^2\delta \tag{10}$$

where:

- $\beta = \sum_{m=1}^{M}\beta_m$: linear "regression" coefficient of the genetic value on the parent A genome proportion
- $\delta = \sum_{m=1}^{M-1}\sum_{m'>m}^{M}\delta_{mm'}$: quadratic "regression" coefficient of the genetic value on the parent A genome proportion, which drives directional epistasis

#### Covariance between genetic values

Let $\mathrm{Cov}(G_i, G_j|\pi_{ik}, \pi_{jk}, \theta_{ij}^k)$ be the covariance between genetic values conditional on shared proportion of alleles originating from each parent:

$$\mathrm{Cov}(G_i, G_j|\pi_{ik}, \pi_{jk}, \theta_{ij}^k) = \Delta_{ij}\sigma_{S_{ij}}^2 + \Delta_{ij}^2\sigma_{S\times S}^2$$

where:

- $\sigma_{S_{ij}}^2 = \sum_{m=1}^{M}\beta_{im}\beta_{jm}$ is the segregation variance generated by substituting parent B alleles by parent A alleles at loci over the whole genome. Note that $\sigma_{S_{ij}}^2$ is specific of the pair of individuals $i$ and $j$, but is practically estimated as an overall parameter $\sigma_S^2$ as it generally varies only a little except when the contribution of epistasis to the trait is very high.
- $\beta_{im} = \left(\beta_{mA} + \sum_{k=1}^{2}\pi_{ik}\sum_{m'\neq m}^{M}\delta_{mm'Ak}\right) - \left(\beta_{mB} + \sum_{k=1}^{2}\pi_{ik}\sum_{m'\neq m}^{M}\delta_{mm'Bk}\right)$: mean effect of substituting parent B alleles by parent A alleles at locus $m$ in a genetic background with $\pi_{ik}$ parent proportions.
- $\sigma_{S\times S}^2 = \sum_{m=1}^{M-1}\sum_{m'>m}^{M}\left(\delta_{mm'}\right)^2$: segregation $\times$ segregation ($S\times S$) interaction variance generated by the simultaneous substitution of parent B alleles by parent A alleles at loci pairs.

#### Derivation of the covariance

An equivalent expression of the model in Eq. (1) is:

$$G_i = \mu + \pi_{iA}\beta + \pi_{iA}^2\delta + \sum_{m=1}^{M}A_{im}^*\beta_{im} + \sum_{m=1}^{M-1}\sum_{m'>m}^{M}A_{im}^*A_{im}^*\delta_{mm'} \tag{11}$$

where:

- $A^*_{im} = A_{imA} - \pi_{iA}$: allele ancestry centered by parent $A$ genome proportion for individual $i$ at locus $m$
- $\text{Cov}\left(A^*_{im}, A^*_{jm}\right) = \Delta_{ij} \ \forall i, j, m$
- $\text{Cov}\left(A^*_{im}A^*_{im'}, A^*_{jm}A^*_{jm'}\right) = \Delta^2_{ij} \ \forall i, j, m \neq m'$
- $\text{Cov}\left(A^*_{im}, A^*_{jm'}\right)^* \approx 0 \ \forall i, j, m \neq m'$
- $\text{Cov}\left(A^*_{im}, A^*_{jm}A^*_{jm'}\right) \approx 0 \ \forall i, j, m \neq m'$
- $\text{Cov}\left(A^*_{im}A^*_{im'}, A^*_{jm}A^*_{jm''}\right) \approx 0 \ \forall i, j, m \neq m', m \neq m'', m' \neq m''$
- $\text{Cov}\left(A^*_{im}A^*_{im'}, A^*_{jm''}A^*_{jm'''}\right) \approx 0 \ \forall i, j, m \neq m', m \neq m'', m \neq m''', m' \neq m'', m' \neq m''', m'' \neq m'''$

One can simply compute the covariance as following:

$$\begin{aligned}
\text{Cov}(G_i, G_j | \pi_{ik}, \pi_{jk}, \theta^k_{ij}) &= \text{Cov}\Bigg(\mu + \pi_{iA}\beta + \pi^2_{iA}\delta + \sum_{m=1}^{M} A^*_{im}\beta_{im} + \sum_{m=1}^{M-1}\sum_{m'>m}^{M} A^*_{im}A^*_{im}\delta_{mm'}, \\
&\qquad \mu + \pi_{jA}\beta + \pi^2_{jA}\delta + \sum_{m=1}^{M} A^*_{jm}\beta_{jm} + \sum_{m=1}^{M-1}\sum_{m'>m}^{M} A^*_{jm}A^*_{jm}\delta_{mm'}\Bigg) \\
\text{Cov}(G_i, G_j | \pi_{ik}, \pi_{jk}, \theta^k_{ij}) &= \sum_{m=1}^{M} \text{Cov}(A^*_{im}, A^*_{jm})\beta_{im}\beta_{jm} + \sum_{m=1}^{M-1}\sum_{m'>m}^{M} \text{Cov}\left(A^*_{im}A^*_{im'}, A_{jm}A^*_{jm'}\right)\left(\delta_{mm'}\right)^2 \\
\text{Cov}(G_i, G_j | \pi_{ik}, \pi_{jk}, \theta^k_{ij}) &= \Delta_{ij}\sigma^2_{S_{ij}} + \Delta^2_{ij}\sigma^2_{S\times S}
\end{aligned}$$
