## Supplementary material for "Uncovering directional epistasis in bi-parental populations using genomic data": File S2

### File S2: Precision of parameter estimates, false positive rate and statistical power of epistasic tests

The precision of parameter estimates as well as the false positive rate and the statistical power of epistasic tests were evaluated with simulated traits using the American maize NAM genotypic data. Only the 24 populations with at least 180 progenies were considered, i.e. excluding the B73×Ki3 population.

#### Simulated traits

Genetic values were simulated using the following genetic model also presented in Eq. (1) from File S1 and equivalent to Eq. (1) in the main text:

$$G_i = \sum_{k \in S} \sum_{m=1}^{M} A_{imk}\beta_{mk} + \sum_{k \in S} \sum_{k' \in S} \sum_{m=1}^{M-1} \sum_{m'>m}^{M} A_{imk}A_{im'k'}\delta_{mm'kk'} \tag{1}$$

where $G_i$ is the genetic value of individual $i$, $S = \{A, B\}$ is the set of parents, $M$ is the number of QTL, $A_{imk}$ is the allele ancestry indicator taking the value 1 if individual $i$ is homozygous for parent $k$ alleles at locus $m$, and 0 otherwise, $\beta_{mk}$ is the main QTL allele effect associated with a homozygous genotype for parent $k$ alleles at locus $m$, and $\delta_{mm'kk'}$ is the deviation QTL allele effects associated with the homozygous genotype for parent $k$ alleles at locus $m$ and for parent $k'$ at locus $m'$. Note that this model was preferred to the equivalent model in Eq. (1) of the main text as the way the alleles effects are defined simplifies the calculation of the simulated segregation variance $\sigma_S^2$, which is used to assess the precision of its estimate (see next section below).

Two genetic scenarios were defined regarding QTL allele effects and are summarized in Table A: "No-epistasis" refers to a trait with only main QTL allele effects and "Epistasis" includes both main and interaction QTL allele effects. A hundred loci were sampled among all SNPs to be used as QTLs. Allele effects were sampled independently from normal distributions with variances defined by the genetic scenario. In the scenario "Epistasis", the number of effects drawn was $2M$ for $\beta_{mk}$ and $4\frac{M(M-1)}{2}$ for $\delta_{mm'kk'}$, which supported the use of $\sigma_\delta^2 = \frac{1}{M-1}$ to obtain a comparable contribution of the main and epistatic effects to the genetic variance. The directionality of epistasis was not directly controlled but was generated by the randomness of simulated QTL effects. Note that even if the expectation of the distribution of $\delta_{mm'kk'}$ is zero, the resulting sum $\delta = \sum_{m=1}^{M-1} \sum_{m'>m}^{M} \delta_{mm'}$, with $\delta_{mm'} = \delta_{mm'kAA} + \delta_{mm'BB} - \delta_{mm'AB} - \delta_{mm'BA}$, has some variance and can substantially deviate from 0, which generates directional epistasis.

Table A: Variance of the normal distribution for allele effects sampled according to the scenario: "No-epistasis" or "Epistasis". The variance $\sigma_\beta^2$ corresponds to the variance of allele effects $\beta_{mk}$, and the variance $\sigma_\delta^2$ corresponds to the variance of allele effects $\delta_{mm'kk'}$

| **Genetic scenario** | $\sigma_\beta^2$ | $\sigma_\delta^2$ |
|---|---|---|
| No-epistasis | 1 | 0 |
| Epistasis | 1 | $\frac{1}{M-1}$ |

The genetic value of each individual was computed as the sum of QTL effects conditional on allele ancestries. Error terms were sampled independently from a normal distribution $\mathcal{N}(0, \sigma_E^2)$ with

$\sigma_E^2$ chosen to reach a given heritability $h^2 = \frac{\sigma_G^2}{\sigma_G^2+\sigma_E^2}$, with $\sigma_G^2 = \Delta_{ii}\sigma_S^2 + \Delta_{ii}^2\sigma_{S\times S}^2$ being the total genetic variance and $\Delta_{ii} = \pi_{iA}\pi_{iB} = \pi_i(1-\pi_i)$ (i.e. the diagonal elements of the $\boldsymbol{\Delta}$ matrix ) with $\pi_i = 0.5$ chosen as the parental proportions of an average individuals. The error covariance matrix $\boldsymbol{D}$ considered in the model was the same as in the real data (see Table 1 from the main text).

The impact of heritability on the precision of variance estimates, the false positive rate and power of statistical tests was evaluated by simulating three contrasted scenarios: $h^2 \in \{0.25, 0.5, 0.75\}$. Similarly the impact of population size was investigated by considering random samples of progenies of size $N \in \{60, 90, 120, 150, 180\}$.

#### Assessment of the precision of parameter estimates

The fixed parameters ($\beta$ and $\delta$) and variance components ($\sigma_S^2$ and $\sigma_{S\times S}^2$) estimates were compared to their reference value calculated using the QTL allele effects of the simulated trait. Note that the exact expression of $\sigma_S^2$ (i.e. $\sigma_{S_{ij}}^2$, see File S1 for details) involves parental proportions $\pi_{ik}$ specific to each individual. The reference value was computed by assuming equal parental proportions in the progeny (i.e., $\pi_{iA} = \pi_{iB} = \frac{1}{2}\ \forall i$). Because of the standardization applied to covariance matrices for the estimation (see Material and Methods), the reference $\sigma_S^2$ was multiplied by $\pi_{iA}\pi_{iB} = \frac{1}{4}$ and the reference $\sigma_{S\times S}^2$ by $(\pi_{iA}\pi_{iB})^2 = \frac{1}{16}$ to make reference and estimated values comparable.

This procedure was replicated 100 times (i.e., over 100 simulated traits), in order to evaluate the mean bias, standard deviation (SD) and root mean square error (RMSE) of the estimates. For a given parameter $\theta$, the mean bias is computed as: $\frac{1}{100}\sum_{i=1}^{100} B_i$ with $B_i = \frac{1}{24}\sum_{j=1}^{24}\left(\widehat{\theta}_{ij} - \theta_i\right)$, $i$ is the $i^{th}$ simulated trait and $j$ is the $j^{th}$ NAM population, the mean SD is computed as: $\frac{1}{100}\sum_{i=1}^{100} S_i$ with $S_i = \sqrt{\frac{1}{24}\sum_{j=1}^{24}\left(\widehat{\theta}_{ij} - \theta_{i.}\right)^2}$, $\theta_{i.}$ is the mean estimate over NAM populations for trait $i$, and the RMSE is computed as: $\frac{1}{100}\sum_{i=1}^{100}\sqrt{B_i^2 + S_i^2}$.

Results are presented in details for both scenarios (i.e., No-epistasis and Epistasis) considering a heritability $h^2 = 0.75$ and a population size $N = 180$ in Table B. As expected, in the "No-epistasis" genetic scenario, the reference epistatic parameters (i.e., $\delta$ and $\sigma_{S\times S}^2$) have null values. For the simulated trait 1, the parameters were generally well estimated in both genetic scenarios. This procedure was replicated over 100 simulated traits to study the mean bias, SD and RMSE. In general parameters showed limited RMSE over all simulated traits.

Table B: Mean and SD of parameter estimates for a given simulated trait (Simulated trait 1) over the 24 American NAM populations according to the genetic scenario (No-epistasis and Epistasis) with $h^2 = 0.75$ and $N = 180$. Mean bias, SD and RMSE of parameter estimates are presented over 100 replicates (Simulated traits 1 to 100)

| | | **Simulated trait 1** | | | **Simulated traits 1 to 100** | | |
|---|---|---|---|---|---|---|---|
| | Parameter | Reference | Mean | SD | Mean bias | Mean SD | Mean RMSE |
| No-epistasis | $\beta$ | -5.51 | -5.79 | 2.58 | -0.06 | 2.12 | 2.34 |
| No-epistasis | $\delta$ | 0.00 | 0.86 | 3.84 | 0.10 | 2.92 | 3.10 |
| No-epistasis | $\sigma_S^2$ | 49.40 | 40.17 | 5.88 | -5.89 | 6.83 | 11.54 |
| No-epistasis | $\sigma_{S\times S}^2$ | 0.00 | 2.06 | 2.59 | 2.62 | 3.80 | 4.86 |
| Epistasis | $\beta$ | -8.36 | -7.98 | 3.47 | -0.13 | 3.09 | 3.81 |
| Epistasis | $\delta$ | 15.68 | 15.14 | 4.85 | 0.14 | 4.07 | 4.67 |
| Epistasis | $\sigma_S^2$ | 73.41 | 71.85 | 11.58 | -9.14 | 12.62 | 20.47 |
| Epistasis | $\sigma_{S\times S}^2$ | 12.40 | 13.38 | 7.94 | -0.72 | 11.40 | 11.92 |

For other heritabilities and population sizes, mean bias, SD and RMSE of parameter estimates are presented over 100 replicates in Table D. In summary, the RMSE of parameters tends to increase with decreasing population size and heritability, which can largely be attributed to a higher SD. In addition to the increase in SD, the positive bias observed for $\sigma^2_{S\times S}$ estimates tends to increase with decreasing population size and heritability.

#### False positive rate

The false positive rate of both epistatic tests was investigated based on the "No-epistasis" genetic scenario. In this genetic scenario both $\sigma^2_{S\times S}$ and $\delta$ take null values. For each population size and heritability, the false positive rate was evaluated as the proportion of significant tests ($\alpha = 5\%$) among the combinations of 24 populations and 100 simulated traits. For both tests, the observed proportion of significant tests was generally consistent with the expected proportion, with a maximum of 6.667% for the directional epistasis test when $N = 60$ and $h^2 = 0.75$ and a maximum of 5.792% for the epistatic variance test when $N = 120$ and $h^2 = 0.50$ (Table C).

Table C: False positive rate of epistatic tests according to population size ($N$) and heritability ($h^2$) calculated as the proportion of significant tests ($\alpha = 5\%$) among 100 traits simulated using the "No-epistasis" genetic scenario and the 24 American maize NAM populations. Tests include (i) the directional epistasis test ($\delta$), and (ii) the epistatic variance test ($\sigma^2_{S\times S}$).

| $N$ | $h^2$ | $\delta$ | $\sigma^2_{S\times S}$ |
|---|---|---|---|
| 60 | 0.25 | 6.625% | 3.750% |
| 60 | 0.50 | 6.500% | 3.208% |
| 60 | 0.75 | 6.667% | 1.167% |
| 90 | 0.25 | 5.583% | 4.958% |
| 90 | 0.50 | 5,125% | 5.625% |
| 90 | 0.75 | 6.417% | 3.583% |
| 120 | 0.25 | 5.625% | 5.250% |
| 120 | 0.50 | 5.417% | 5.792% |
| 120 | 0.75 | 5.667% | 4.708% |
| 150 | 0.25 | 5.542% | 4.875% |
| 150 | 0.50 | 5.292% | 5.667% |
| 150 | 0.75 | 5.208% | 5.750% |
| 180 | 0.25 | 5.625% | 4.917% |
| 180 | 0.50 | 5.625% | 4.917% |
| 180 | 0.75 | 5.292% | 5.542% |

#### Statistical power

The statistical power of both epistatic tests was investigated based on the "Epistasis" genetic scenario.

In this genetic scenario, $\sigma^2_{S\times S}$ takes non-null values and vary only a little from a simulated trait to another (mean of 12.50 and a standard deviation of 0.25). For each population size and heritability, the statistical power was defined as the proportion of significant tests ($\alpha = 5\%$) among the combinations of 24 populations and 100 simulated traits. The statistical power obtained on

Table D: Mean bias, SD and RMSE of parameter estimates over the 24 American NAM populations and 100 simulated traits according to the genetic scenario (No-epistasis and Epistasis), the population size $N$ and the heritability $h^2$

| | | | $\beta$ | | | $\delta$ | | | $\sigma^2_S$ | | | $\sigma^2_{S\times S}$ | | |
|---|---|---|---|---|---|---|---|---|---|---|---|---|---|---|
| Scenario | $N$ | $h^2$ | Bias | SD | RMSE | Bias | SD | RMSE | Bias | SD | RMSE | Bias | SD | RMSE |
| Epistasis | 60 | 0.25 | 0.467 | 10.882 | 11.785 | -0.377 | 13.663 | 14.452 | -10.225 | 61.666 | 69.043 | 58.833 | 95.423 | 118.197 |
| Epistasis | 60 | 0.50 | -0.080 | 7.276 | 7.897 | 0.212 | 8.754 | 9.303 | -13.115 | 35.572 | 43.257 | 19.633 | 39.551 | 45.994 |
| Epistasis | 60 | 0.75 | -0.456 | 5.382 | 6.005 | 0.490 | 6.073 | 6.635 | -12.381 | 24.874 | 32.624 | 3.076 | 18.292 | 19.441 |
| Epistasis | 90 | 0.25 | -0.585 | 9.102 | 9.940 | 0.282 | 12.132 | 12.823 | -10.265 | 47.055 | 54.250 | 48.330 | 83.476 | 101.856 |
| Epistasis | 90 | 0.50 | -0.442 | 6.061 | 6.735 | 0.442 | 7.634 | 8.177 | -10.770 | 27.876 | 35.164 | 14.250 | 33.091 | 37.716 |
| Epistasis | 90 | 0.75 | -0.161 | 4.208 | 4.807 | 0.183 | 4.953 | 5.447 | -10.644 | 19.127 | 26.868 | 2.030 | 16.255 | 17.054 |
| Epistasis | 120 | 0.25 | -0.319 | 8.299 | 9.478 | 0.453 | 11.417 | 12.324 | -10.045 | 38.747 | 46.404 | 36.342 | 67.208 | 81.006 |
| Epistasis | 120 | 0.50 | -0.458 | 5.349 | 6.200 | 0.463 | 7.092 | 7.748 | -10.033 | 22.449 | 30.258 | 9.767 | 28.318 | 31.728 |
| Epistasis | 120 | 0.75 | -0.165 | 3.713 | 4.459 | 0.145 | 4.706 | 5.306 | -9.493 | 15.465 | 23.550 | 0.831 | 14.325 | 14.928 |
| Epistasis | 150 | 0.25 | -0.804 | 7.711 | 8.751 | 0.636 | 11.086 | 11.855 | -8.539 | 34.668 | 41.916 | 27.175 | 54.989 | 65.208 |
| Epistasis | 150 | 0.50 | -0.342 | 4.905 | 5.651 | 0.357 | 6.731 | 7.304 | -9.281 | 19.815 | 27.160 | 7.417 | 25.213 | 27.585 |
| Epistasis | 150 | 0.75 | -0.298 | 3.320 | 4.134 | 0.202 | 4.328 | 4.951 | -9.446 | 13.665 | 21.794 | 0.235 | 12.739 | 13.276 |
| Epistasis | 180 | 0.25 | -0.173 | 7.310 | 8.409 | 0.174 | 10.772 | 11.599 | -9.298 | 29.882 | 37.349 | 20.576 | 45.446 | 52.659 |
| Epistasis | 180 | 0.50 | -0.251 | 4.551 | 5.385 | 0.063 | 6.436 | 7.090 | -9.007 | 17.861 | 25.565 | 4.487 | 20.706 | 22.601 |
| Epistasis | 180 | 0.75 | -0.130 | 3.087 | 3.810 | 0.137 | 4.073 | 4.674 | -9.135 | 12.618 | 20.472 | -0.722 | 11.404 | 11.921 |
| No-epistasis | 60 | 0.25 | 0.427 | 7.940 | 8.673 | -0.244 | 10.124 | 10.701 | -5.305 | 35.476 | 39.918 | 38.914 | 51.574 | 67.162 |
| No-epistasis | 60 | 0.50 | 0.203 | 5.369 | 5.764 | -0.169 | 6.452 | 6.818 | -7.591 | 20.588 | 25.247 | 15.246 | 19.205 | 25.497 |
| No-epistasis | 60 | 0.75 | 0.148 | 3.658 | 3.907 | -0.085 | 4.215 | 4.431 | -7.299 | 13.378 | 18.322 | 5.698 | 7.418 | 9.783 |
| No-epistasis | 90 | 0.25 | 0.142 | 6.893 | 7.490 | -0.251 | 9.134 | 9.631 | -6.480 | 27.409 | 32.327 | 28.049 | 41.087 | 52.040 |
| No-epistasis | 90 | 0.50 | -0.135 | 4.343 | 4.613 | -0.005 | 5.576 | 5.812 | -6.398 | 15.311 | 19.904 | 11.868 | 16.400 | 21.285 |
| No-epistasis | 90 | 0.75 | 0.050 | 3.035 | 3.227 | -0.033 | 3.695 | 3.872 | -6.701 | 10.090 | 15.250 | 4.779 | 6.168 | 8.149 |
| No-epistasis | 120 | 0.25 | 0.088 | 8.515 | 9.107 | 0.025 | 5.154 | 5.432 | -6.521 | 21.861 | 26.016 | 22.578 | 34.565 | 43.971 |
| No-epistasis | 120 | 0.50 | -0.185 | 3.869 | 4.197 | 0.205 | 5.154 | 5.432 | -6.250 | 12.527 | 17.515 | 9.148 | 13.088 | 16.903 |
| No-epistasis | 120 | 0.75 | 0.020 | 2.554 | 2.814 | 0.021 | 3.244 | 3.450 | -6.367 | 8.398 | 13.355 | 3.681 | 5.096 | 6.609 |
| No-epistasis | 150 | 0.25 | -0.081 | 5.690 | 6.530 | -0.094 | 8.222 | 8.755 | -6.351 | 19.260 | 24.146 | 18.663 | 27.794 | 35.446 |
| No-epistasis | 150 | 0.50 | -0.037 | 3.576 | 3.982 | 0.069 | 4.992 | 5.316 | -5.764 | 11.268 | 16.061 | 7.396 | 11.077 | 14.213 |
| No-epistasis | 150 | 0.75 | -0.124 | 2.318 | 2.519 | 0.093 | 3.085 | 3.261 | -6.166 | 7.342 | 12.372 | 3.173 | 4.523 | 5.806 |
| No-epistasis | 180 | 0.25 | -0.242 | 5.544 | 6.237 | 0.246 | 8.267 | 8.794 | -5.530 | 17.142 | 22.032 | 16.076 | 23.993 | 31.194 |
| No-epistasis | 180 | 0.50 | 0.010 | 3.375 | 3.789 | -0.003 | 4.863 | 5.170 | -5.802 | 10.258 | 14.928 | 5.805 | 8.606 | 11.059 |
| No-epistasis | 180 | 0.75 | -0.058 | 2.117 | 2.339 | 0.101 | 2.923 | 3.100 | -5.894 | 6.834 | 11.541 | 2.620 | 3.803 | 4.864 |

those simulations was generally very low, except for large populations with high $h^2$ where the power was moderate with a maximum of 19.000% (Table E).

For $\delta$, the absolute simulated value may be close to null or large depending on the sampling (mean of -4.30 and a standard deviation of 13.86 over the 100 simulated traits). In this context, the statistical power was also evaluated as the proportion of significant tests ($\alpha = 5\%$), but considering simulated traits either globally or grouped into quartiles according to the absolute value of the simulated $\delta$. Statistical power was close to the type I error of $\alpha = 5\%$ for the lowest $|\delta|$ interval and could increase close to 1 for the largest $|\delta|$ interval (Table E). It also increased with increasing population size and heritability.

In summary, the directional epistasis test is generally more statistically powerful than the epistatic variance test.

Table E: Statistical power of the two epistatic test procedure according to population size ($N$) and heritability ($h^2$) calculated as the proportion of significant tests ($\alpha = 5\%$) among 100 traits simulated using the "Epistasis" genetic scenario and the 24 American maize NAM populations. Tests include (i) the directional epistasis test ($\delta$), and (ii) the epistatic variance test ($\sigma^2_{S\times S}$). For $\delta$, simulated traits were considered either globally (All) or clustered into quartiles according to the absolute value of the simulated $\delta$: the first quartile with $|\delta| \in [0.040, 3.48[$, the second quartile with $|\delta| \in [3.48, 7.96[$, the third quartile with $|\delta| \in [7.96, 13.12[$ and the last quartile with $|\delta| \in [13.12, 34.68[$.

| | | | $\delta$ | | | | |
|---|---|---|---|---|---|---|---|
| $N$ | $h^2$ | $\sigma^2_{S\times S}$ | All | 1st quartile | 2nd quartile | 3rd quartile | 4th quartile |
| 60 | 0.25 | 4.542% | 16.292% | 7.667% | 9.167% | 15.167% | 31.944% |
| 60 | 0.50 | 4.583% | 27.083% | 7.333% | 13.833% | 26.667% | 59.028% |
| 60 | 0.75 | 3.250% | 39.042% | 8.833% | 20.000% | 44.500% | 82.118% |
| 90 | 0.25 | 6.875% | 16.000% | 5.000% | 7.667% | 15.333% | 34.375% |
| 90 | 0.50 | 6.917% | 28.917% | 6.333% | 11.500% | 29.500% | 67.014% |
| 90 | 0.75 | 8.667% | 42.333% | 6.167% | 20.333% | 53.833% | 88.542% |
| 120 | 0.25 | 6.708% | 18.375% | 6.000% | 10.500% | 16.167% | 39.236% |
| 120 | 0.50 | 8.000% | 31.208% | 5.667% | 15.833% | 31.167% | 71.007% |
| 120 | 0.75 | 12.042% | 46.875% | 9.333% | 29.333% | 56.000% | 92.535% |
| 150 | 0.25 | 7.167% | 18.083% | 3.667% | 8.167% | 17.667% | 41.667% |
| 150 | 0.50 | 10.208% | 33.083% | 5.333% | 13.667% | 36.167% | 76.215% |
| 150 | 0.75 | 15.500% | 46.750% | 5.500% | 26.500% | 60.000% | 94.792% |
| 180 | 0.25 | 6.792% | 20.417% | 6.000% | 12.333% | 13.333% | 48.611% |
| 180 | 0.50 | 9.292% | 32.417% | 7.333% | 14.667% | 32.500% | 74.132% |
| 180 | 0.75 | 19.000% | 48.417% | 6.500% | 26.333% | 64.833% | 95.833% |
