## Supplementary material for "Uncovering directional epistasis in bi-parental populations using genomic data": Figure S1, S2, S3, S4, S5, S6, S7 and S8

### Detecting directional and non-directional epistasis in bi-parental populations using genomic data

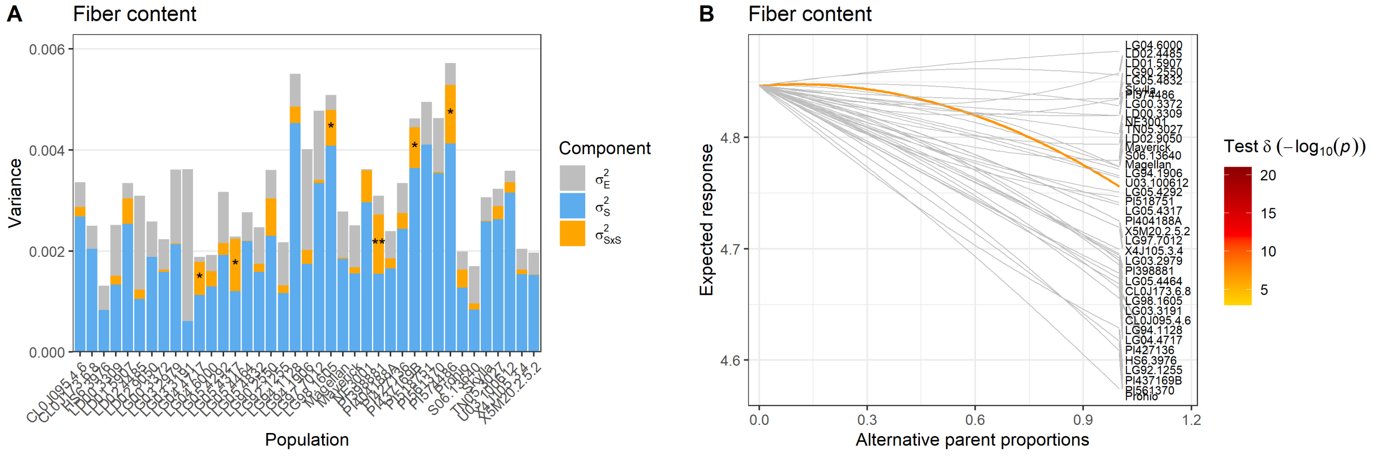


**Figure S1**: Variance component barplots (**A**) and directional epistasis plots (**B**) of the soybean NAM for fiber content. For the variance component barplot, the range of p-value obtained for the likelihood ratio test of $\sigma^2_{S\times S}$ are indicated as: *** for p-values inferior to 0.001, ** for p-values inferior to 0.01, and * for p-values inferior to 0.05. For the directional epistasis plot, tests with a p-value higher than a Bonferroni threshold of 5% over the number of populations are indicated in grey.

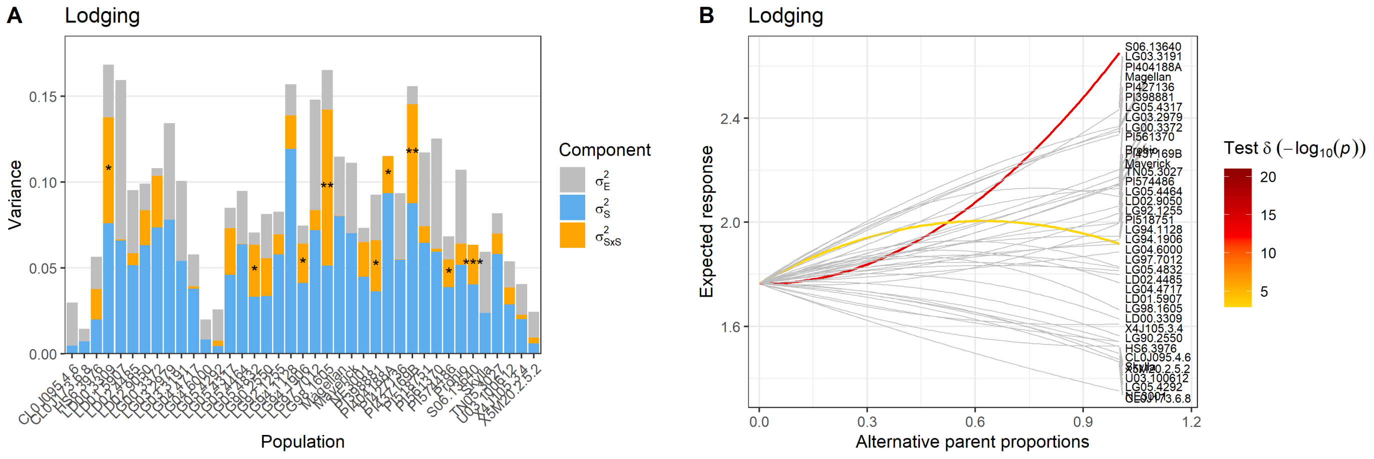


**Figure S2**: Variance component barplots (**A**) and directional epistasis plots (**B**) of the soybean NAM for lodging. For the variance component barplot, the range of p-value obtained for the likelihood ratio test of $\sigma^2_{S\times S}$ are indicated as: *** for p-values inferior to 0.001, ** for p-values inferior to 0.01, and * for p-values inferior to 0.05. For the directional epistasis plot, tests with a p-value higher than a Bonferroni threshold of 5% over the number of populations are indicated in grey.

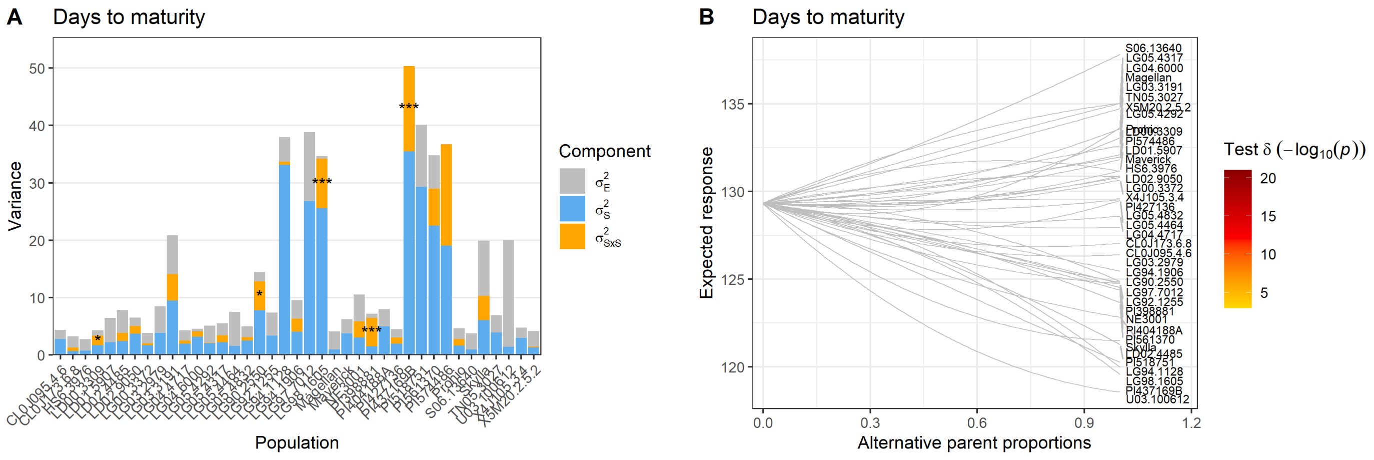


**Figure S3**: Variance component barplots (**A**) and directional epistasis plots (**B**) of the soybean NAM for days to maturity. For the variance component barplot, the range of p-value obtained for the likelihood ratio test of $\sigma^2_{S\times S}$ are indicated as: *** for p-values inferior to 0.001, ** for p-values inferior to 0.01, and * for p-values inferior to 0.05. For the directional epistasis plot, tests with a p-value higher than a Bonferroni threshold of 5% over the number of populations are indicated in grey.

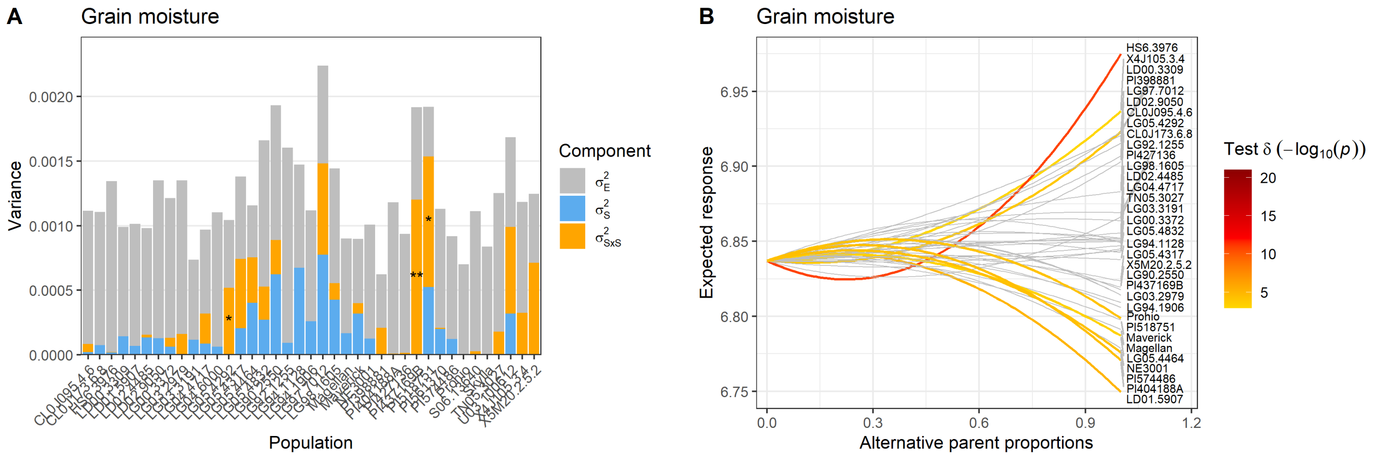


**Figure S4**: Variance component barplots (**A**) and directional epistasis plots (**B**) of the soybean NAM for grain moisture. For the variance component barplot, the range of p-value obtained for the likelihood ratio test of $\sigma^2_{S\times S}$ are indicated as: *** for p-values inferior to 0.001, ** for p-values inferior to 0.01, and * for p-values inferior to 0.05. For the directional epistasis plot, tests with a p-value higher than a Bonferroni threshold of 5% over the number of populations are indicated in grey.

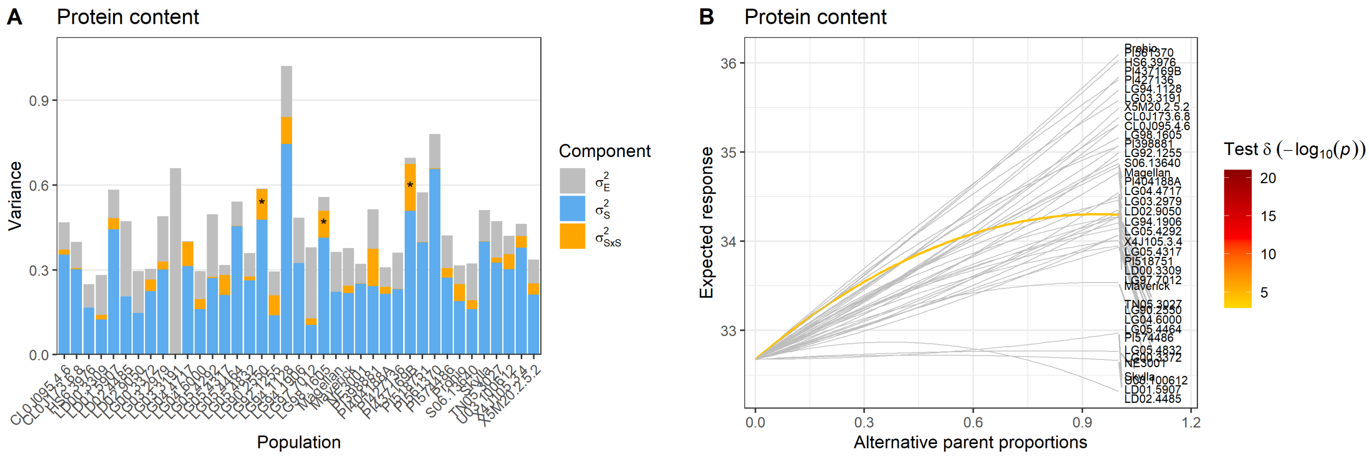


**Figure S5**: Variance component barplots (**A**) and directional epistasis plots (**B**) of the soybean NAM for protein content. For the variance component barplot, the range of p-value obtained for the likelihood ratio test of $\sigma^2_{S\times S}$ are indicated as: *** for p-values inferior to 0.001, ** for p-values inferior to 0.01, and * for p-values inferior to 0.05. For the directional epistasis plot, tests with a p-value higher than a Bonferroni threshold of 5% over the number of populations are indicated in grey.

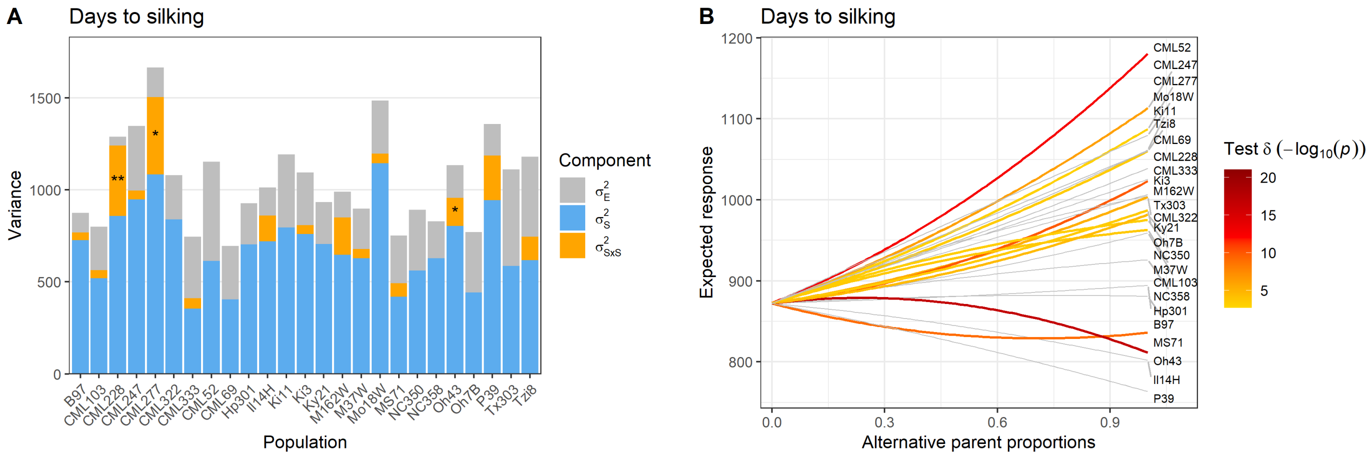


**Figure S6**: Variance component barplots (**A**) and directional epistasis plots (**B**) of the maize NAM for days to silking. For the variance component barplot, the range of p-value obtained for the likelihood ratio test of $\sigma^2_{S\times S}$ are indicated as: *** for p-values inferior to 0.001, ** for p-values inferior to 0.01, and * for p-values higher than 0.05. For the directional epistasis plot, tests with a p-value inferior to a Bonferroni threshold of 5% over the number of populations are indicated in grey.

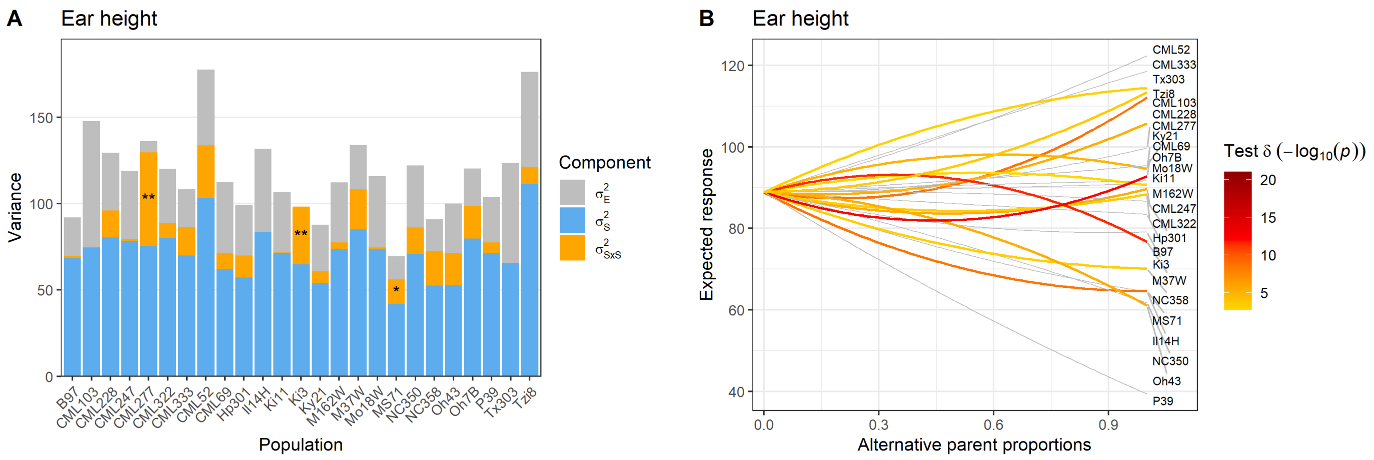


**Figure S7**: Variance component barplots (**A**) and directional epistasis plots (**B**) of the maize NAM for ear height. For the variance component barplot, the range of p-value obtained for the likelihood ratio test of $\sigma^2_{S\times S}$ are indicated as: *** for p-values inferior to 0.001, ** for p-values higher than 0.01, and * for p-values inferior to 0.05. For the directional epistasis plot, tests with a p-value inferior to a Bonferroni threshold of 5% over the number of populations are indicated in grey.

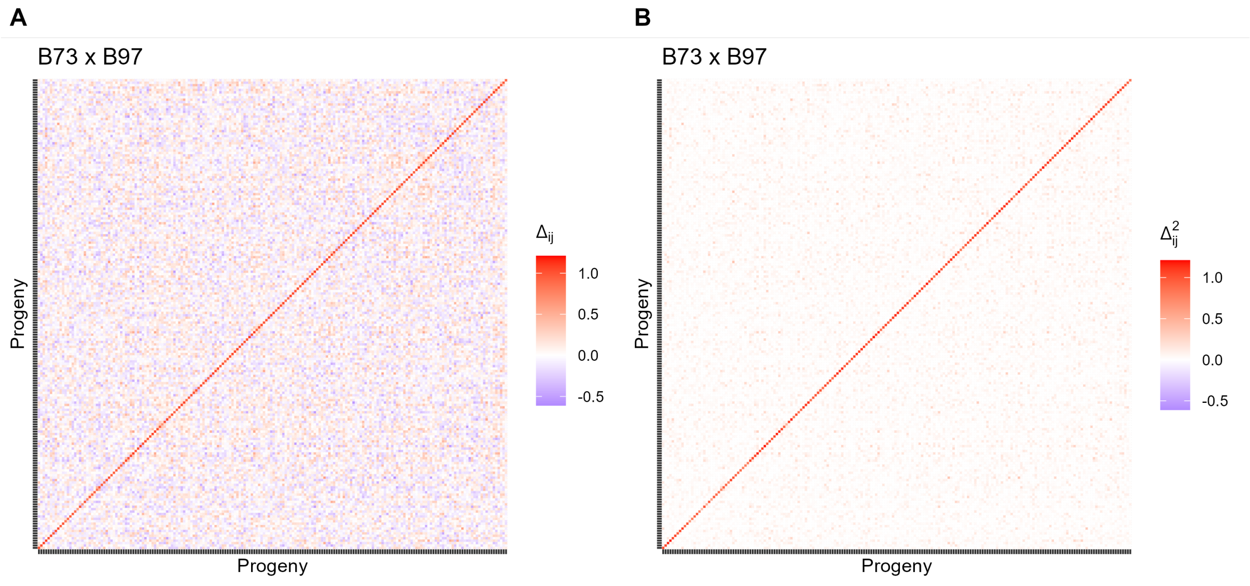


**Figure S8**: Coefficients of (**A**) $\Delta_{ij}$ and (**B**) $\Delta^2_{ij}$ for the maize bi-parental population B73×B97 after the standardization of the $\mathbf{\Delta}$ and $\mathbf{\Delta}^2$ matrices (see materials and methods for details). For $\Delta_{ij}$, the diagonal elements take values in the interval $[0.62, 1.10]$ and off-diagonal elements take values in the interval $[-0.61, 0.61]$. For $\Delta^2_{ij}$, the diagonal elements take values in the interval $[0.38, 1.21]$ and off-diagonal elements take values in the interval $[0.00, 0.37]$
